# Long-read sequencing of extrachromosomal circular DNA and genome assembly of a *Solanum lycopersicum* breeding line revealed active LTR retrotransposons originating from *S. peruvianum* L. introgressions

**DOI:** 10.1101/2023.12.18.572100

**Authors:** Pavel Merkulov, Melania Serganova, Georgy Petrov, Vladislav Mityukov, Ilya Kirov

## Abstract

Transposable elements (TEs) are a major force in the evolution of plant genomes. Differences in the transposition activities and landscapes of TEs can vary substantially, even in closely related species. Interspecific hybridization, a widely employed technique in tomato breeding, results in the creation of novel combinations of TEs from distinct species. The implications of this process for TE transposition activity have not been studied in modern cultivars. In this study, we used nanopore sequencing of extrachromosomal circular DNA (eccDNA) and identified two highly active Ty1/*Copia* LTR retrotransposon families of tomato (*Solanum lycopersicum*), called *Salsa* and *Ketchup*. Elements of these families produce thousands of eccDNAs under controlled conditions and epigenetic stress. EccDNA sequence analysis revealed that the major parts of eccDNA produced by *Ketchup* and *Salsa* exhibited low similarity to the *S. lycopersicum* genomic sequence. To trace the origin of these TEs, whole-genome nanopore sequencing and de novo genome assembly were performed. We found that these TEs occurred in a tomato breeding line via interspecific introgression from *S. peruvianum*. Our findings collectively show that interspecific introgressions can contribute to both genetic and phenotypic diversity not only by introducing novel genetic variants, but also by importing active transposable elements from other species.

## 1. Introduction

Transposable elements are ubiquitous components of plant genomes. LTR retrotransposons (LTR-RTEs) are among the highest copy members of the plant mobilome (Feschotte, Jiang, & Wessler, 2002; Zemach et al., 2013), accounting for 76% of the rye genome (G. Li et al., 2021) and 50% of the tomato genome (Su et al., 2021). Uncontrolled TE reactivation can lead to genetic instability. Therefore, plants have a complex system of epigenetic control of TEs, including RNA-dependent DNA Methylation (RdDM) (Cuerda-Gil & Slotkin, 2016; Matzke, Kanno, & Matzke, 2015) and *DDM1*-mediated DNA methylation (Zemach et al., 2013). TEs in plants carrying mutations in the genes involved in epigenetic regulation can be transcriptionally reactivated. For example, a threefold increase in TE transcription was observed in a triple mutant of *A. thaliana* with mutations in three key TE-controlling genes, *ddm1*, rdr6, and polV (Panda & Slotkin, 2020). Similarly, transposition of some TE families has been observed in the DNA methylation-free *A. thaliana* mutant (He et al., 2022), which is likely due to the redistribution of histone modifications (Zhao et al., 2022).

TE transposition can also occur in wild type plants under natural conditions. Such natural TE activity has contributed significantly to the evolution, adaptation, and domestication of plants (Lisch, 2013). Bursts in LTR retrotransposon activity have had a key impact on the size of the genomes of some plant species (Kim et al., 2017; Penin et al., 2021; Vitte & Panaud, 2005), whereas individual insertions have created functionally altered or new alleles of genes (Cai et al., 2022; Galindo-Gonzalez, Mhiri, Deyholos, & Grandbastien, 2017). More specific examples include the emergence of traits such as seedless apples (Yao, Dong, & Morris, 2001), grape skin color (Kobayashi, Goto-Yamamoto, & Hirochika, 2004), and red orange pulp (Butelli et al., 2012). In tomatoes, the insertion of elements of the *Rider* family resulted in an elongated fruit shape (N. Jiang, Gao, Xiao, & van der Knaap, 2009; Xiao, Jiang, Schaffner, Stockinger, & van der Knaap, 2008), yellow flesh fruit (N. V. Jiang, S.; Wu, S.; Knaap E Van Der, 2012), and lack of formation of a detachment zone in the peduncle (Roldan et al., 2017). A study on *A. thaliana* also provided an interesting example of the contribution of TE insertion to plant adaptation to a novel ecological environment. A rare variant of the *A. thaliana* FLC locus was found to contain an intronic insertion of a heat-inducible element in the *ONSEN* family, which may be an adaptation to flowering in the absence of vernalization (Quadrana, 2020). Whole genome sequencing of *A. thaliana* ecotypes revealed that hundreds of TEs generate novel insertions (Quadrana, 2020). Despite this, the exact triggers of TE transposition activity are known only for a small set of elements, such as the heat-induced *ONSEN* LTR retrotransposon of *A. thaliana* and the plant tissue culture-triggered Tos17 LTR retrotransposon of rice (Dubin, Mittelsten Scheid, & Becker, 2018).

In addition to environmental factors, TE activity can be triggered by ‘genomic shock’ as proposed by McClintock (McClintock, 1984). Genomic shock can result from chromosomal rearrangements and interspecific hybridization. Transcriptional reactivation of LTR-RTE has been detected during long-distance hybridization in various plant species, including rice (Liu & Wendel, 2000), wheat (Kashkush, Feldman, & Levy, 2002), *Arabidopsis* (Madlung et al., 2005), and wild potato species (Paz, Rendina Gonzalez, Ferrer, & Masuelli, 2015). In addition, although much less frequently, hybridization-induced LTR-RTE transposition in real time has been detected in rice (H. Y. Wang et al., 2010), poplar (Usai et al., 2020), and potato (Gantuz et al., 2022). The consequences of interspecific hybridization on TE composition in the genome have been well described for the *Solanum* genus. Interspecific hybridization is one of the main sources of genetic diversity in tomato (*Solanum lycopersicum* L.) breeding and domestication. A wide range of wild species has been implicated in this process, including *S. peruvianum* (Kaloshian et al., 1998; Tanksley et al., 1998), S. chilense (Zamir et al., 1994), and *S. habrochaites* (X. Yang et al., 2014), *S. penelli* (Bolger et al., 2014) and *S. pimpinellifolium* (C. Zhang et al., 2014). A recent study demonstrated that the TE composition of modern tomato cultivars is less divergent than that of wild species (Dominguez et al., 2020). At the same time, the domesticated tomato *S. lycopersicum* var. cerasiforme shows minor losses in the number of mobile TE families (Dominguez et al., 2020), which could potentially be due to recurrent hybridization with its closest wild relative, *S. pimpinellifolium* (Blanca et al., 2015). Whether TEs located in interspecific introgressions maintain their activity during breeding and generate new insertions in the recipient genome has not been well studied.

In this study, we aimed to decipher inducible mobilome activity originating from TEs located at interspecific introgressions in the tomato genome. We performed a whole-genome analysis of TE activity using long-read nanopore sequencing of extrachromosomal circular DNAs from a tomato line. We found thousands of eccDNAs mapped on members of families that we called ‘*Ketchup*’ and ‘*Salsa*.’ The eccDNA sequence analysis revealed that the major parts of eccDNA produced by *Ketchup* and *Salsa* did not fully align with any tomato (SL3.0) reference TEs, but were similar to the TEs from the *S. peruvianum* genome. We performed whole-genome nanopore sequencing and assembly for our tomato line and revealed large interspecific introgressions carrying members of the *Ketchup* and *Salsa* TE families. Our results suggested that active TEs introduced by interspecific hybridization may serve as an additional source of genetic diversity during plant breeding.

## 2. Results

### 2.1. Mobilome-seq of a tomato breeding line

To unravel TEs capable of completing their life cycle, we performed nanopore (ONT) sequencing of extrachromosomal circular DNA (eccDNAs) of a tomato plant. To collect more active TEs, we grew the plants in a special medium containing a mixture of zebularine and α-amanitin (A&Z). These chemicals lead to DNA methylation reduction and inhibition of Polymerase II, a major player in the PolII-RDR6 TE RdDM silencing pathway (Thieme et al., 2017). Plants grown in MS medium without A&Z were used as controls to statistically evaluate eccDNA peaks.

**Figure 1.**
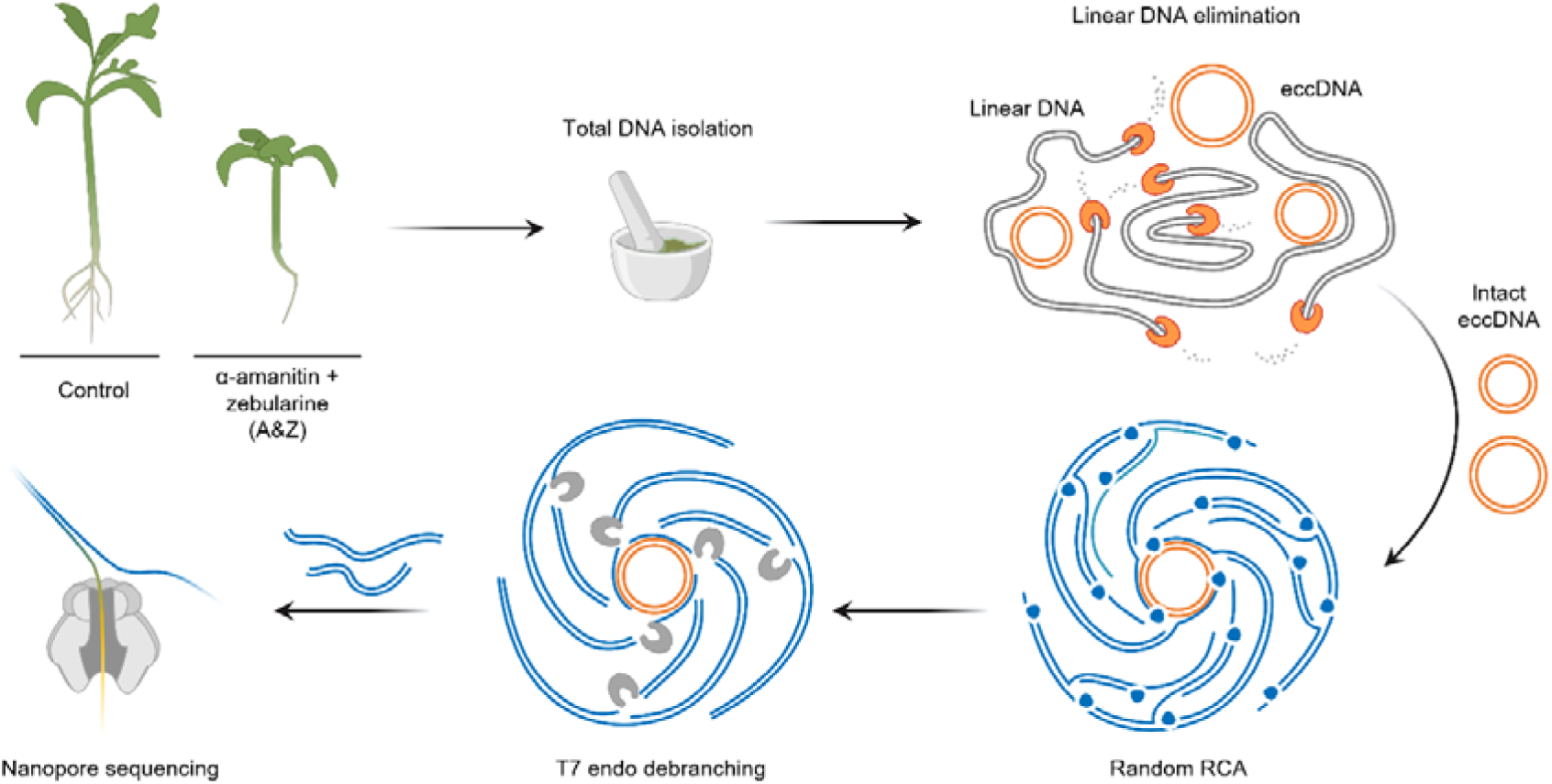
Overview of the eccDNA nanopore sequencing experiment.

In total, we obtained 64,000 and 48,000 reads for the A&Z and Control plants, respectively. The eccDNA reads were mapped to the reference genome (SL3.0), followed by intersection with LTR-RTE coordinates (4206 annotated LTR-RTEs) and manual curation. We found 101 LTR-RTEs, for which >10 ONT reads were accounted. Of these, 38 LTR-RTEs demonstrated significant overrepresentation of ONT reads from the A&Z sample compared with the control sample (Fisher’s exact test with multiple correction p-value < 0.01). Phylogenetic analysis of these LTR-RTEs based on their LTR sequences revealed that they belong to two families, that we named ‘*Salsa*’ (a tomato-based sauce that dates back to the earliest tomato cultivators, the Aztecs and Mayans (Ibsen & Nielsen, 1999)) and ‘*Ketchup*’ (Figure S1). According to the GyDB database classification, the *Ketchup* elements belong to the Tork clade and are almost identical to *CopiaSL_35* (average 98% identity with 99% coverage), whereas the *Salsa* family belongs to the Bianca clade and has some similarity to *CopiaSL_25* (average about 72% identity with 41 % coverage). Among the elements of each family, we selected one RTE with the highest CPM value: RTE976 (Sly11:3,646,824..3,652,356) from *Salsa* and RTE511 (Sly01:68,869,427..68,874,507) from *Ketchup* family. We checked for the presence of open reading frames in the genomic sequences of selected RTEs in the SL3.0 genome assembly, and both elements were found to have one or two long ORFs (Figure S2A).

**Figure 2.**
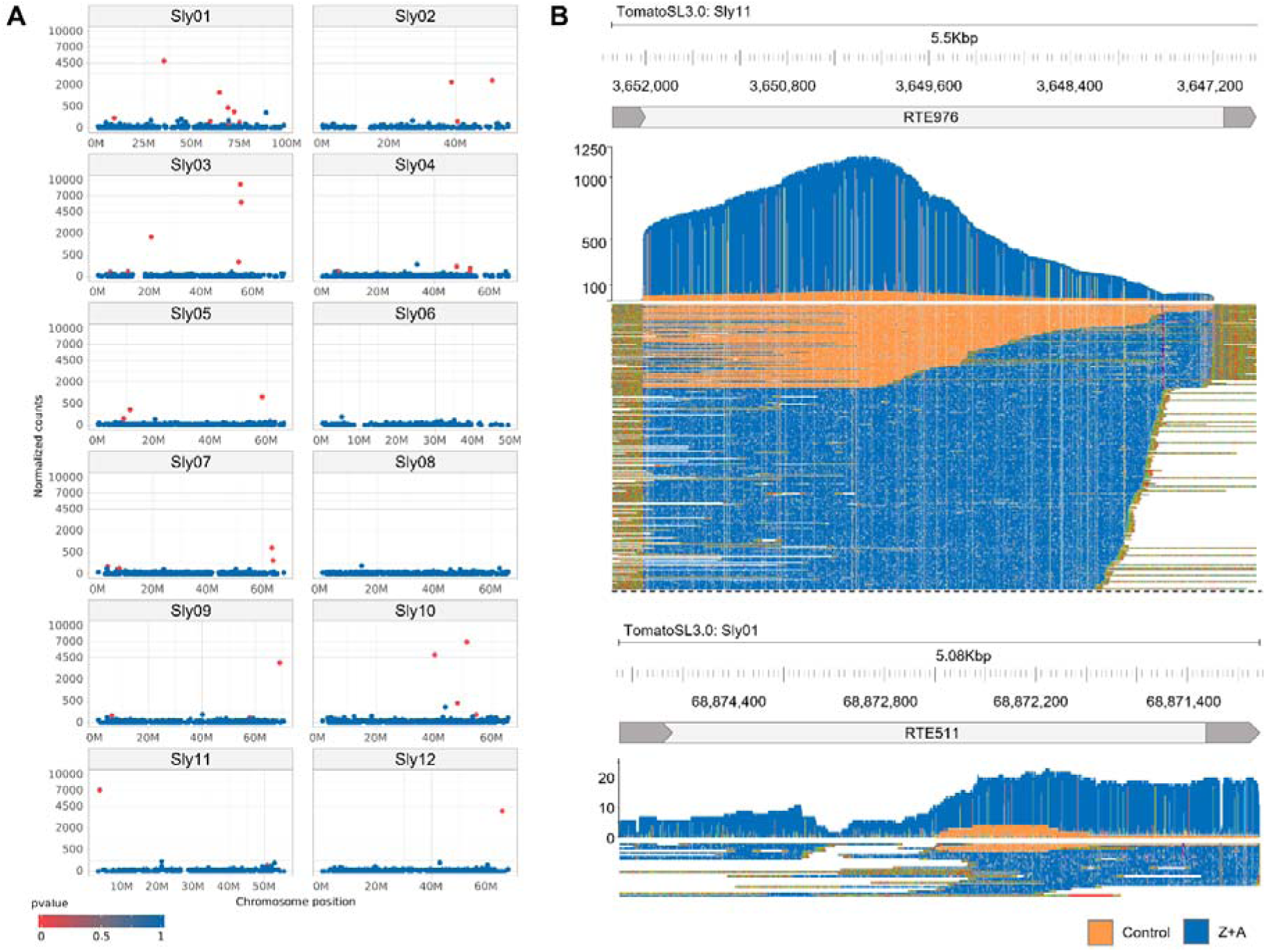
Analysis of eccDNA production by LTR-RTEs assessed using ONT sequencing. (A) Normalized (reads per 100,000 ONT reads) eccDNA count for 4206 RTEs located on chromosomes of SL3.0 genome assembly; (B) Coverage of RTE976 (*Salsa* family) and RTE511 (*Ketchup* family) by eccDNA reads. Orange and blue colors indicate eccDNA reads from the Control and Z&A samples.

Thus, nanopore Mobilome-Seq revealed that under epigenetic stress conditions (A&Z treatment), tens of LTR-RTEs belonged to two distinct families of tomato lines producing eccDNAs.

### 2.2 RTEs producing eccDNAs originate from *S. peruvianum*

Surprisingly, however, a detailed analysis of LTR-RTE coverage by ONT eccDNA reads revealed numerous SNPs (>50 for RTE511 and >100 for RTE976), distinguishing the LTR-RTE sequences of our tomato line from the reference. In addition, LTRs of RTE976 were not covered by eccDNA reads, and LTRs of RTE511 possessed a >20bp deletion based on read mapping. These observations show that the eccDNA reads most probably originated from RTEs that were not present in the reference genome sequence (SL3.0). To verify this, we performed a BLAST search for the most similar LTR sequences in the genomic assemblies of wild relatives of *S. lycopersicum*, including *S. penelli*, *S. lycopersicum* var. cerasiforme, *S. pimpinellifolium*, S. lycopersicoides, *S. peruvianum*, S. chilense, *S. habrochaites*. We used consensus LTR sequences deduced from the eccDNA reads mapped to RTE976/RTE511 as queries for the BLAST search. This analysis revealed that the most similar sequences were found in the *S. peruvianum* (SP) genome with >96% identity to the query LTR sequences (Figure 3A).

**Figure 3.**
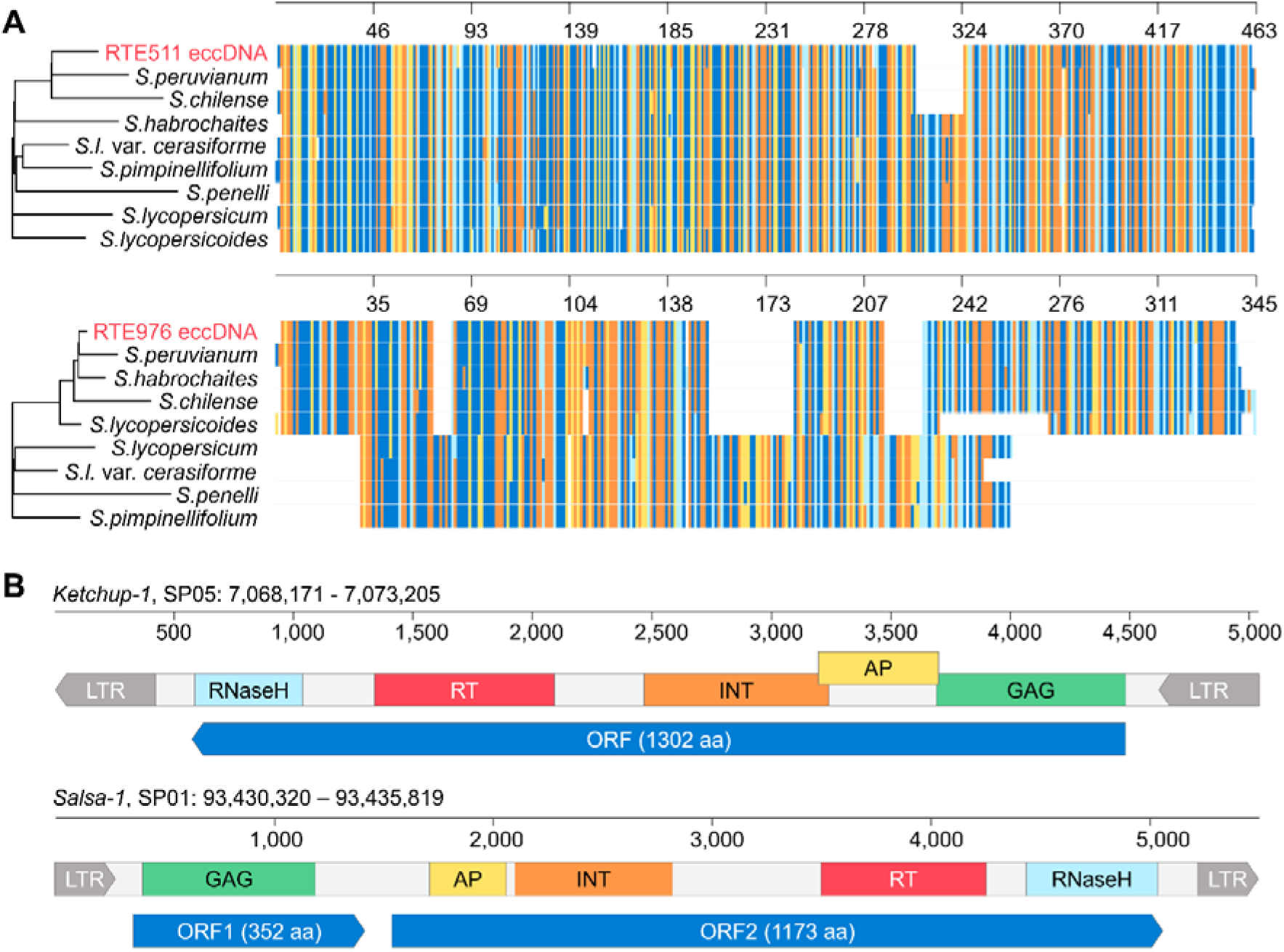
Phylogenetic and structural analyses of *Ketchup* and *Salsa* RTEs. (A) Alignment of LTR sequences from eccDNA and the genome assemblies of diverse tomato species. (B) Structure and open reading frames of two RTEs (*Ketchup-1* and *Salsa-1*) in the *S. peruvianum* genome.

We identified the full-length RTEs (5033 and 5500 bp) of SP genome with LTRs that have >98% similarity to eccDNA deduced LTRs and named them *Ketchup-1* (SP05:7,068,171-7,073,205) and *Salsa-1* (SP01:93,430,320-93,435,819). Both RTEs had 99-100% LTR identity, suggesting their recent activity in the SP genome. Both RTEs possessed well-defined reading frames (1302 aa for *Ketchup-1*; 352 and 1173 aa for *Salsa-1*) encoding all the required domains, including GAG coat protein (GAG), aspartic proteinase (AP), integrase (INT), reverse transcriptase () and ribonuclease H (RNaseH) (Figure 3B).

### 2.3 *Ketchup-1* and *Salsa-1* produce full-length eccDNAs with one or two LTRs

Individual RTE eccDNAs may represent different structural variants covering only a small LTR part, as well as whole RTEs (Merkulov, Egorova, & Kirov, 2023). To assess the structure of eccDNAs produced by *Ketchup-1* and *Salsa-1*, we investigated the monomers of individual eccDNA reads possessing concatemers. We found that a significant portion of eccDNAs of *Ketchup-1* contained only one LTR, with some eccDNAs also possessing small deletions (Figure 4A). In turn, *Salsa-1* produced full-length eccDNA with one or more LTR (Figure 4B) and a small proportion of eccDNAs containing truncated sequences (Figure S3). We then performed inverted PCR using specific primers for amplification of the LTR junction regions of eccDNA (Figure 4C). For this experiment, genomic DNA from Control and A&Z samples before and after eccDNA enrichment and RCA were used. Weak and strong PCR products were obtained for solo-LTR eccDNAs produced by *Ketchup-1* and *Salsa-1* in control and Z&A samples, respectively (Figure 4C). However, we were unable to detect extrachromosomal linear DNA (eclDNA) for *Salsa-1* and *Ketchup-1* in either the control or Z&A samples (Figure S4), suggesting that these RTEs are not capable of producing insertions under laboratory conditions.

**Figure 4.**
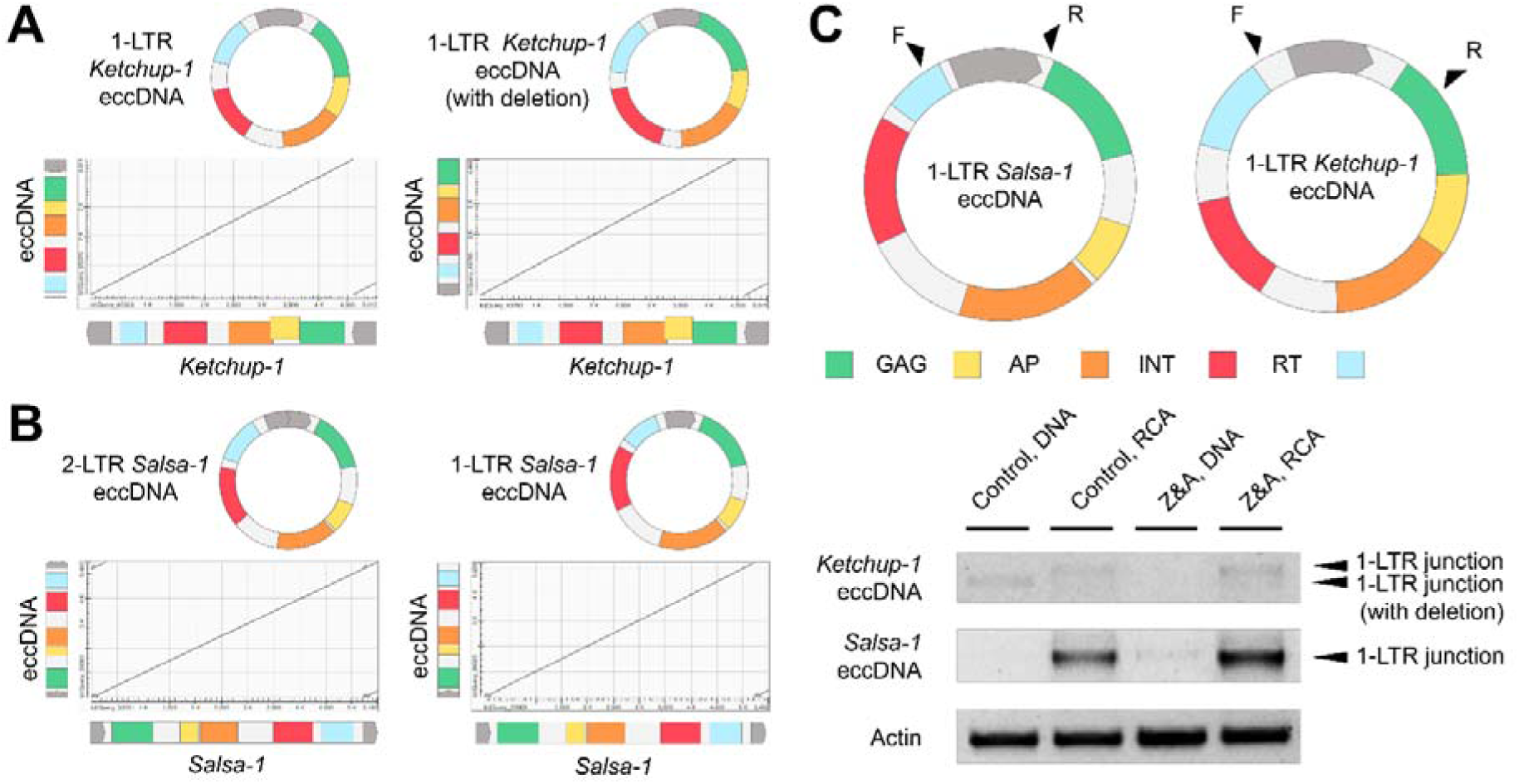
Analysis of the structure of *Salsa-1* and *Ketchup-1* eccDNA. Dot plot from alignment of eccDNA deduced monomer sequences against full-length reference *Ketchup-1* (A) and *Salsa-1* sequences. (C) Primer positions (top) and gel electrophoresis (bottom) for inverted PCR with genomic DNA from Control and A&Z samples before and after eccDNA enrichment and RCA.

Altogether, the results demonstrate that *Ketchup-1* and *Salsa-1* RTEs produce eccDNAs with one or two LTRs under control and A&Z conditions.

### 2.4 *Ketchup-1* and *Salsa-1* invaded *S. lycopersicum* genome via an interspecific introgression

To determine how SP RTEs occurred in the genome of our tomato line, we performed whole-genome nanopore sequencing. We obtained 4,417,781 reads with a total length of 57.7Gb corresponding to ∼64x genome coverage of the tomato genome (1C=900Mb, (Tomato Genome, 2012)). We performed SNP calling and found significant biases in SNP density along SL chromosomes. The results unambiguously demonstrated a significantly high density of SNPs along full-length chromosomes 6 (2Mb–32Mb) and 9 (2Mb–64Mb) (Figure 5A; Figure S5). These results indicated that the genome of the tomato line used in this study possessed large interspecific introgressions.

**Figure 5.**
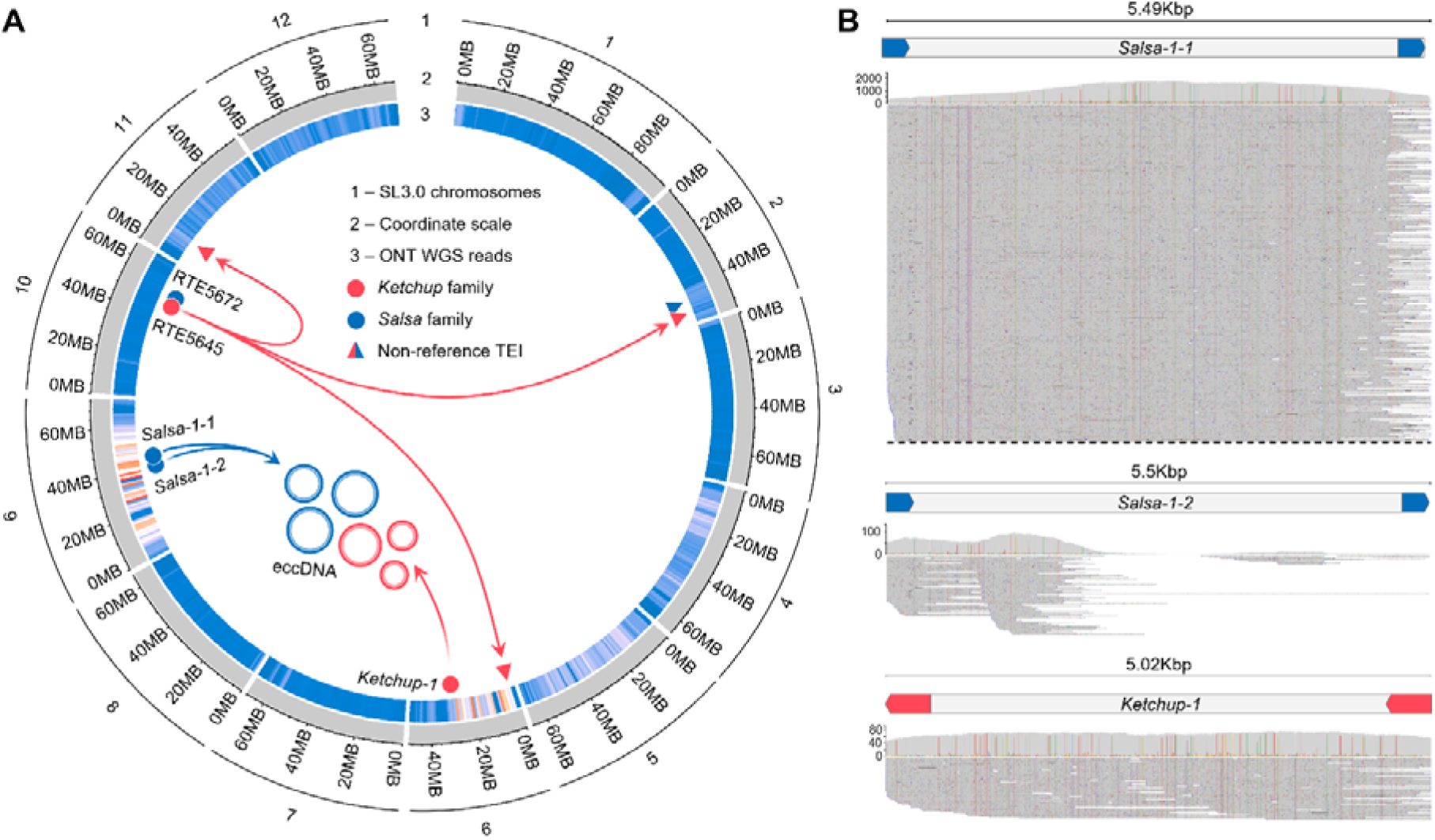
Whole-genome nanopore sequencing of the analyzed tomato line. (A) SNP density deduced from alignment of ONT WGS reads of the studied tomato line on SL3.0 genome assembly; circles and triangles indicate original TEs and their insertions; rings represent eccDNAs produced by *Ketchup* and *Salsa* of *S. peruvianum*. (B) Mapped eccDNA reads from Z&A samples on contigs from ONT data.

To gain further insight into the origin of *Salsa* and *Ketchup* in our tomato line, we performed whole-genome assembly using only WGS nanopore reads. The assembly was performed using NextDenovo (Hu et al., 2023). The draft assembly resulted in an N50 around 16.7 Mb (16.721.217 bp) and total length of approximately 800Mb (813056715 bp). The quality of the draft assembly was verified using the BUSCO software (Manni, Berkeley, Seppey, & Zdobnov, 2021). The percentage of the BUSCOs benchmark genes was high (>98%). A comparison of the assembled genome and reference SL3.0 revealed that SL chromosomes 2, 4, and 7 were almost completely covered by two assembled contigs, further suggesting the relatively high contiguity of the draft genome (Figure S5). In line with the SNP density distribution, the comparison also revealed a low alignment rate between our assembly and chromosomes SL6 and SL9, pointing to the genomic differences between SL3.0 and the genome of our breeding line. Comparing our draft genome assembly with SL3.0 showed the position of *Salsa* and *Ketchup* SP retrotransposon insertions on SL3.0 reference: *Salsa-1-1* (Sly9:46,243,555) and *Salsa-1-2* (Sly09:42,506,399), as well as a single insertion of *Ketchup-1*, localized on chromosome 6 (Sly06:31,315,480).

We also identified three and one new insertions for the *Ketchup* and *Salsa* RTEs of *S. lycopersicum*, RTE5645 (TEIs: Sly02, 51,094,794; Sly06, 2,425,515; and Sly11, 9,441,876) and RTE5672 (TEIs: Sly02:46,224,366), respectively. All TEIs were validated using PCRs with primers targeting the flanking regions and TEs (Figure S6A and S6B). Using the TEI junctions, we checked for the presence of insertions at the same sites among the genomic assemblies: *S. penelli*, *S. cerasiforme*, *S. pimpinellifolium*, S. lycopersicoides, *S. peruvianum*, S. chilense, *S. habrochaites*, *S. galapagense*, *S. neorickii*, *S. chmielewskii*, *S. lycopersicum* cv. M82, and *S. lycopersicum* cv. ZY65. Only the RTE5672 insertion was detected in selected genomic assemblies: *S. pimpinellifolium* (2:38,850,750-38,856,308) and *S. galapagense* (2:46,113,449-46,119,003), indicating the recent occurrence of other non-reference insertions.

Thus, using WGS ONT data, we showed that *Salsa-1* and *Ketchup-1* occurred in the genome of our tomato line via interspecific hybridization and chromosomal introgression.

## 3. Discussion

Interspecific hybridization has been extensively used to introduce desirable genes from wild species into cultivated tomato (Blanca et al., 2015). It has been known for a long time that interspecific hybridization may trigger TE reactivation and transposition (McClintock, 1984). The TE composition of the tomato genome has been described previously (Dominguez et al., 2020). Here, we explored whether interspecific introgression might bring novel active TEs from other species. Using nanopore sequencing of eccDNAs, we described real-time mobilome activity that occurred under epigenetic stress in a tomato breeding line. The sequences of individual eccDNA reads allowed us to accurately determine two families of active LTR retrotransposons: *Salsa* and *Ketchup*. Further elucidation of the newly obtained draft genome assembly for our breeding line revealed that the eccDNA-producing RTEs from these two families were introgressed from *S. peruvianum*.

Our results highlight how active transposons can be introduced into a new genome to maintain their activity for several generations. Indeed, hybridization-induced TE mutagenesis can be a major factor in the evolution of sexually reproducing organisms (Fukai et al., 2022), and it has even been exploited for crop improvement (Paszkowski, 2015). Interestingly, the population of transpositionally active TEs in wild tomato species is significantly larger than that in cultivated *S. lycopersicum* (Dominguez et al., 2020). Therefore, interspecific chromosomal introgressions in modern tomato varieties may carry active TEs.

Interestingly, we did not observe any novel insertions of introgressed RTEs in the genome of our breeding line. This can be partially explained by the transposition of the original elements in the first stages of hybridization, followed by their subsequent elimination during backcrossing and selection. In contrast to the introgressed SP RTEs, we identified novel insertions for SL members of the *Salsa* and *Ketchup* families. It is interesting to speculate that the presence of active *Salsa* RTEs from *S. peruvianum* complemented SL RTEs to transpose, as has been shown for BARE-2 and Tos17 GAG-defective elements (Sabot, 2014; Tanskanen, Sabot, Vicient, & Schulman, 2007).

Short read sequencing has been frequently used for eccDNA detection (Esposito et al., 2019; Kwolek et al., 2022; Lanciano et al., 2017). Utilization of long-read WGS and eccDNA sequencing allowed us to accurately determine the structure and full-length sequence of the eccDNAs. This allowed us to identify the positions of the active elements that are absent in the reference genome. Although the formation of eccDNA originating from RTEs has been considered a by-product of their activities (Garfinkel et al., 2006), a recent study suggested that eccDNA is one of the key steps in the life cycle of RTEs (F. Yang et al., 2023). The concatameric structure of the eccDNA ONT reads allows distinguishing naturally occurring truncated sequences from DNA breaks that occur during the sample preparation procedure. This feature of eccDNA ONT data can help shed light on the composition and origin of eccDNA in cells (Merkulov et al., 2023; P. Zhang et al., 2023). The authors showed that the *ONSEN* and *EVD* elements produced almost equal amounts of full-length (> 5 Kb) and truncated (< 1000bp) eccDNAs in the *ddm1* background. Interestingly, growing *Arabidopsis* on A&Z medium resulted in a shift in eccDNA composition toward truncated eccDNAs (Merkulov et al., 2023). These results are in contrast with our results for the *Salsa* and *Ketchup* elements. Here, we showed that *Salsa* and *Ketchup* TEs mainly produced full-length eccDNAs with one or two LTRs in tomato plants grown on A&Z media. These results suggest that eccDNA formation under similar growth conditions (for example, A&Z) may differ for different species and TEs.

In addition to the production of eccDNA under the relaxation of epigenetic control, *Salsa* and *Ketchup* also exhibited activity in the control sample, although to a lesser extent. EccDNA production poses a serious threat to genomic integrity and stability. The generation of eccDNA may result in genomic rearrangement via spontaneous reintegration into the genome, as has been shown for various types of eccDNA in eukaryotes (Arrey, Keating, & Regenberg, 2022). In addition, it has been suggested that a high load of eccDNA may alter DNA repair pathways, leading to new genetic variations (P. Zhang et al., 2023). Additionally, eccDNAs may serve as a template for transcription of protein coding or non-coding RNAs further expanding the repertoire of possible consequences for the plant (Peng, Mirouze, & Bucher, 2022). For inheritance to the next generation, eccDNA-mediated genetic changes need to be produced in the plant ‘germ line’ cells, such as meristematic cells of the shoot apical meristem (SAM), pollen, or egg cells. However, RTE transcription and transposition are limited in these cells through specific epigenetic mechanisms (Nguyen & Gutzat, 2022). Thus, it remains an open question whether *Salsa* and *Ketchup* are capable of generating novel genetically inherited insertions, and whether their eccDNAs contribute to genome instability. This question could be answered by genomic analysis of M1 plants, which will be the subject of our future research.

## 4. Conclusion

Using nanopore whole-genome and eccDNA sequencing, we identified two novel families of tomato TEs, *Salsa* and *Ketchup*, that produce eccDNAs under both control and epigenetic stress conditions. We showed that these TEs occur in a tomato breeding line via interspecific introgression from *S. peruvianum*. Collectively, our results demonstrate that interspecific introgression may contribute to genetic and phenotypic diversity not only by providing new genetic variants, but also by bringing new active TEs from other species.

## 5. Materials and Methods

### 5.1. Plant material and in vitro growth conditions

Tomato plants were grown on ½ MS medium supplemented with 4 mg/ml α-amanitin and 8 mg/ml zebularine for two weeks under a long-day photoperiod (16/8).

### 5.2. Total DNA isolation

Total DNA was isolated from two-week-old seedlings using the modified CTAB method described by Pucker (https://www.protocols.io/view/plant-dna-extraction-and-preparation-for-ont-seque-kxygxenmkv8j/v1).

### 5.3. eccDNA isolation and sequencing

For eccDNA isolation we used the techniques described by Lanciano et al. (Lanciano et al., 2017) and Wang et al. (Y. Wang, Wang, & Zhang, 2023) with modifications. Briefly, to remove linear DNA, 1 μg of total DNA was treated with 1 μl (10 U/μl) of PlasmidSafe DNase supplemented with 2 μl of ATP (25 mM) and 5 μl of 10× PlasmidSafe buffer in a volume of 50 μl. The reaction was incubated for 72 hours with additional reagents (0.1 μl enzyme, 0.2 μl ATP, 0.3 μl buffer) was added every 24 h, followed by incubation at 72°C for 30 min. Precipitation of eccDNA was carried out by overnight incubation at −20°C in the presence of 0.1 volume of 3 M sodium acetate (pH 5.2) and 2.5 × volume of absolute ethanol, followed by centrifugation at 12,000 × g for 30 min. The eccDNA pellet was washed with ice-cold 70% ethanol and dissolved in 10 μl of deionized nuclease-free water. For eccDNA amplification using random RCA, 2 µL phi29 polymerase (Thermo Scientific, EP0091), 2 µL 10× phi29 reaction buffer, 5 µL 10 mM dNTP, and 1 µL 500 µM exo-resistant oligo (NpNpNpNpNpSNpSN, where p is phosphodiester and pS is the phosphorothioate group) with the addition of nuclease-free water to a final volume of 20 µL. The reaction was preheated to 95°C for 5 min, ramped to 30°C at a 1% ramp rate on a thermocycler, and incubated for 36 h at 30°C. The enzyme was inactivated by heating the mixture at 65*C for 10 min. For debranching, 500 ng of RCA amplicons were treated with T7 endonuclease 5 μL of 10× reaction buffer and 1 μL of T7 endonuclease I (New England Biolabs, M0302S) in a 50 μL reaction volume. After incubation at 37°C for 15 min, the reaction was stopped immediately and purified by adding an equal volume of chloroform. The Debranched RCA product was precipitated by adding 1/10V 3M sodium acetate (pH 5.2) and absolute ethanol (2.5V), followed by incubation at −80°C for 30 min and centrifugation at 12,000× g for 30 min. The pellet obtained was dissolved in nuclease-free water and used for nanopore sequencing.

### 5.4. Nanopore Library Preparation and Sequencing

For eccDNA sequencing, library preparation was carried out with 500 ng of cDNA using Native Barcoding Expansion 1–12 (Oxford Nanopore Technologies (Oxford, UK), catalog no. EXP-NBD104), and the Ligation Sequencing Kit SQK-LSK109 (Oxford Nanopore Technologies). Sequencing was performed using MinION equipped with an R9.4.1 flow cell.

For whole-genome sequencing, a fraction of short fragments was removed from 9 μg of total DNA using the Short-Read Eliminator Kit XL (PacBio, SKU 102-208-400), according to the manufacturer’s recommendations. The library was prepared with 1 μg of long fragment-enriched DNA using the Ligation Sequencing Kit SQK-LSK109 (Oxford Nanopore Technologies). Sequencing was carried out using PromethION P2 equipped with an R9.4.1 flow cell for 72 hours.

### 5.5. Whole-genome sequencing and *de novo* assembly

The raw Nanopore long reads were assembled into sequence contigs using NextDenovo (version 2.5.2) (Hu et al., 2023) with the following parameters 900Mb of estimated genome size and for assembly minimap option, minimum overlap was set to 5000 bp, and other parameters were set by default. Draft assembly resulted in an N50 of approximately 16.7 Mb (16.721.217 bp) and a total length of approximately 800Mb (813056715 bp), which was verified by BUSCO software (v5.5.0) (Manni et al., 2021) with both eukaryota and *viridiplatae* lineages, as well as both metaeuk and miniprot options, and by FastANI (version 1.33) (Jain, Rodriguez, Phillippy, Konstantinidis, & Aluru, 2018) alignment on *Solanum lycopersicum* reference (genome assembly SL3.0). The percentage of complete BUSCOs ranged from 94.5% with miniprot option and eukaryota lineage; 97.8% with miniprot option and *viridiplantae* lineage; 98.0% with metaeuk option and eukaryota lineage to 99.3% with metaeuk option and *viridiplantae* lineage. SNPs were identified using Clair3 software (Luo et al., 2021) and their chromosome distribution was visualized by pycircos python package (https://github.com/ponnhide/pyCircos).

### 5.6. DNA amplification

Amplification was carried out using a Bio-Rad T100™ thermal cycler (Bio-Rad Laboratories, USA). A 25 µl reaction mixture contained: 1 µl DNA (25 ng), 2.5 µl 10× Taq Turbo buffer, 0.2 µl Hot Start Taq polymerase (5 units/μl), 1 µl 10 pmol of each primer, 0.5 µl dNTP (10 mM) and 18.8 µl nuclease-free water.

### 5.7. Validation of the insertions

For validation, 25 ng of total DNA/ecDNA was amplified using RTE-specific inverted PCR primers (Table S2).

### 5.8. eclDNA intermediates detection

For eclDNA amplification, the Sequence-Independent Retrotransposons Trapping (SIRT) method was used (Griffiths, Catoni, Iwasaki, & Paszkowski, 2018). To form SIRT adaptors, equal volumes of 100 μM of SIRT_adaptor_1 (5′-GTAATACGACTCACTATAGGGCACGCGTCCACGACGGCCCGGGCTCCA-3′), and SIRT_adaptor_2 (5′-PO4-TGGAGCCC-3′) oligos were mixed and incubated at 95°C for 10 min, followed by cooling to room temperature. The ligation mixture was prepared on ice with 300 ng of total DNA using 8 μl of adapters, 1.6 μl of 10× overnight buffer, and 1 μl of T4 ligase (100 U/μl), with nuclease-free water added to the final volume. 16 µl. Ligation was performed at 14°C for 16 h, followed by enzyme inactivation at 65°C for 10 min. The entire reaction volume was purified using 0.5 volumes of AMPure XP SPRI Reagent (Beckman Coulter, A63881) according to the manufacturer’s instructions. DNA eluted in 30 μl was amplified using the adaptor-specific primer AP1 (5’-GTAATACGACTCACTATAGGGC-3’) and the TE-specific primers listed in Table S1. The amplification program consisted of 95°C for 3 min and 35 cycles of 95°C for 10 s, 51°C for 10 s, and 72°C for 1 min. The resulting amplicons were separated on 1.5% agarose gel at 80 V for 60 min.

### 5.9. Bioinformatic analysis of eccDNA sequencing and data visualization

Raw eccDNA nanopore reads were mapped to the SL3.0 genome using the minimap2 software (H. Li, 2018) with the following parameters: -ax map-ont -t 100. The obtained SAM files were converted to BAM format, sorted, and indexed using SAMtools (H. Li et al., 2009). To obtain the eccDNA peaks, the obtained sorted bam files were analyzed using the eccStructONT pipeline, as previously described (Merkulov et al., 2023).

For the evolutionary analysis, the genomes of *S. lycopersicum* var. *lycopersicum* (Heinz1706; ver. SL3.0) and *S. lycopersicum* var. *cerasiforme* (LA1673) were downloaded from https://solgenomics.net/. *S. lycopersicum* var. lycopersicum cv. M82, *S. lycopersicum* var. lycopersicum cv. ZY65, *S. penelli* (LA716), *S. pimpinellifolium* (LA1547), S. lycopersicoides (LA2951), *S. peruvianum* (LA0446), S. corneliomulleri (LA1331), *S. neorickii* (LA0247), *S. chmielewskii* (LA1028), S. chilense (LA1969), *S. habrochaites* (LA1777) and *S. galapagense* (LA0436) genome assemblies were downloaded from http://caastomato.biocloud.net/.

An alignment and tree visualisation were made using ggplot2 (version 3.4.4) (Wickham, 2016), ggtree (version 3.8.2) (Yu, Smith, Zhu, Guan, & Lam, 2017) and ggmsa (version 1.6.0) (Zhou et al., 2022).

## Supplementary Materials

The following supporting information can be downloaded at: www.mdpi.com/xxx/s1, Figure S1: Phylogenetic tree of full-length sequences of 38 RTEs; Figure S2: Encoded domains and open reading frames (ORFs) for RTEs; Figure S3: Dot plots for alignment of truncated eccDNA sequences and full-length retrotransposon sequences with domain annotation; Figure S4: PCR validation of eclDNA accumulation for *S. peruvianum* elements; Figure S5: PCR validation of *Ketchup* and *Salsa* TEIs in the tomato line used in the study; Table S1: Filtered RTEs list; Table S2: Primers used in this study.

## Author Contributions

Conceptualization, I.K. and P.M.; methodology, I.K. and P.M.; software, I.K.; validation, M.S. and G.P.; formal analysis, I.K. and P.M.; investigation, I.K., P.M., M.S., G.P. V.M.; resources, I.K.; data curation, P.M., V.M.; writing—original draft preparation, P.M, I.K.; writing—review and editing, I.K.; visualization, P.M., I.K..; supervision, I.K.; project administration, I.K.; funding acquisition, I.K. All authors have read and agreed to the published version of the manuscript.

## Data Availability Statement

The nanopore data produced for this study are available in the Sequence Read Archive (SRA), NCBI, under Bioproject Accession PRJXXXXX.

## Acknowledgments

The authors thank Maria Logacheva (Skolkovo Institute of Science and Technology) and Alexey Penin (The Institute for Information Transmission Problems) for their help with whole genome nanopore sequencing.

---

**Supplementary Figure S1.**
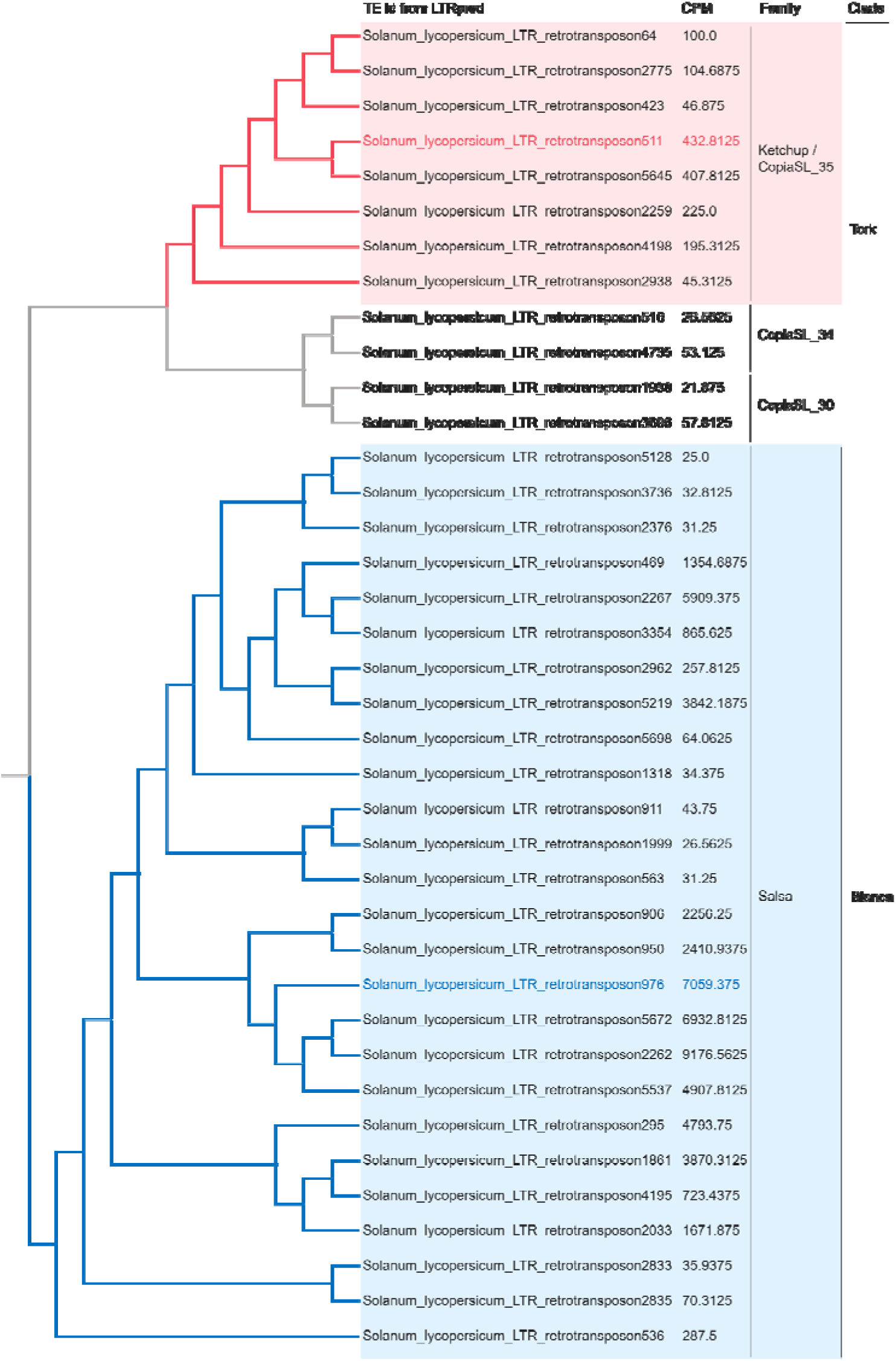
Phylogenetic tree of full-length sequences of 38 RTEs. The red and blue colors correspond to the elements of the *Ketchup* and *Salsa* families, respectively.

**Supplementary Figure S2.**
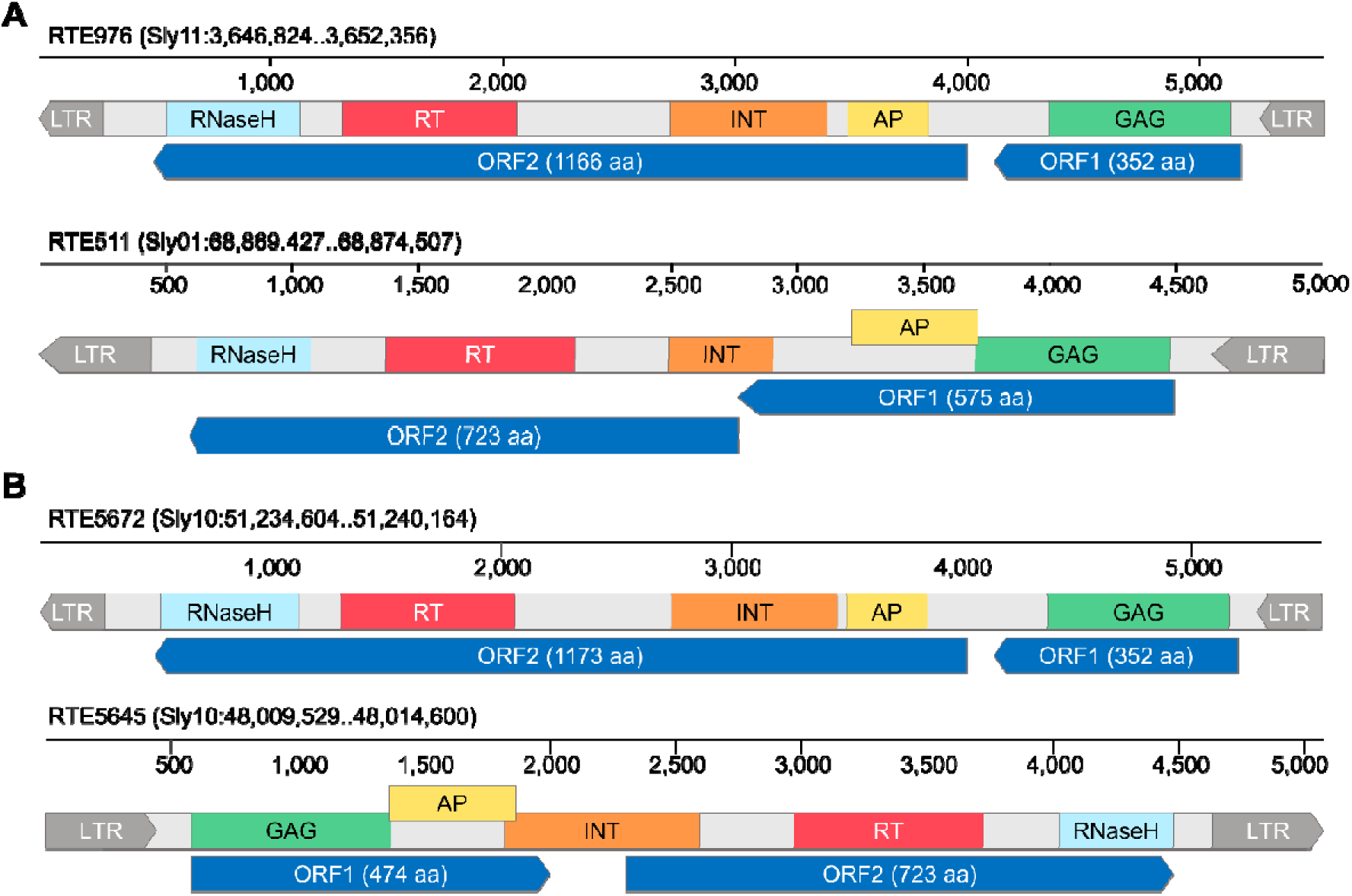
Encoded domains and open reading frames (ORFs) for RTEs (A) RTE976 and RTE511, two SL elements belonging to the *Salsa* and *Ketchup* family respectively; (B) RTE5672 and RTE5645, two SL elements belonging to the *Salsa* and *Ketchup* family respectively.

**Supplementary Figure S3.**
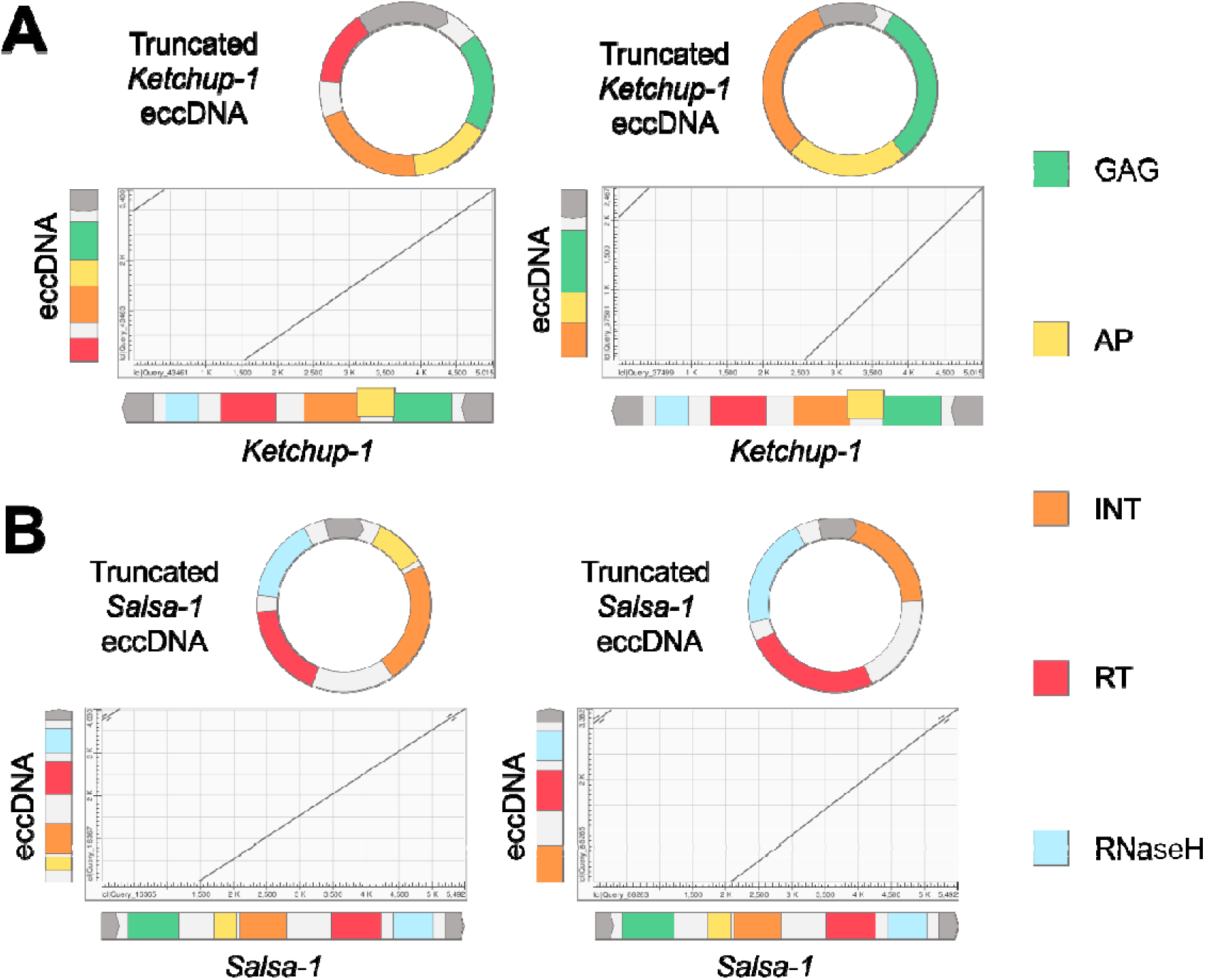
Dot plots for alignment of truncated eccDNA sequences (Y axis) and full-length retrotransposon sequences (X axis) with domain annotation: (A) – *Ketchup-1*; (B) – *Salsa-1*.

**Supplementary Figure S4.**
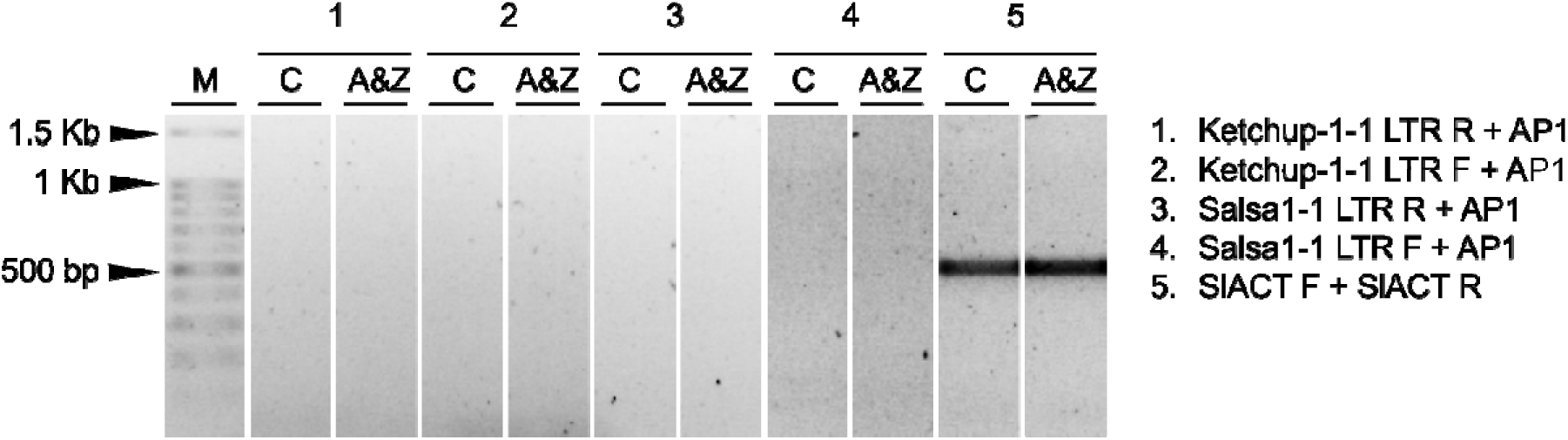
PCR confirmation of the absence of eclDNA accumulation for *Ketchup-1-1* and *Salsa-1-1* elements in control (C) and relaxed TE control (A&Z) samples.

**Supplementary Figure S5.**
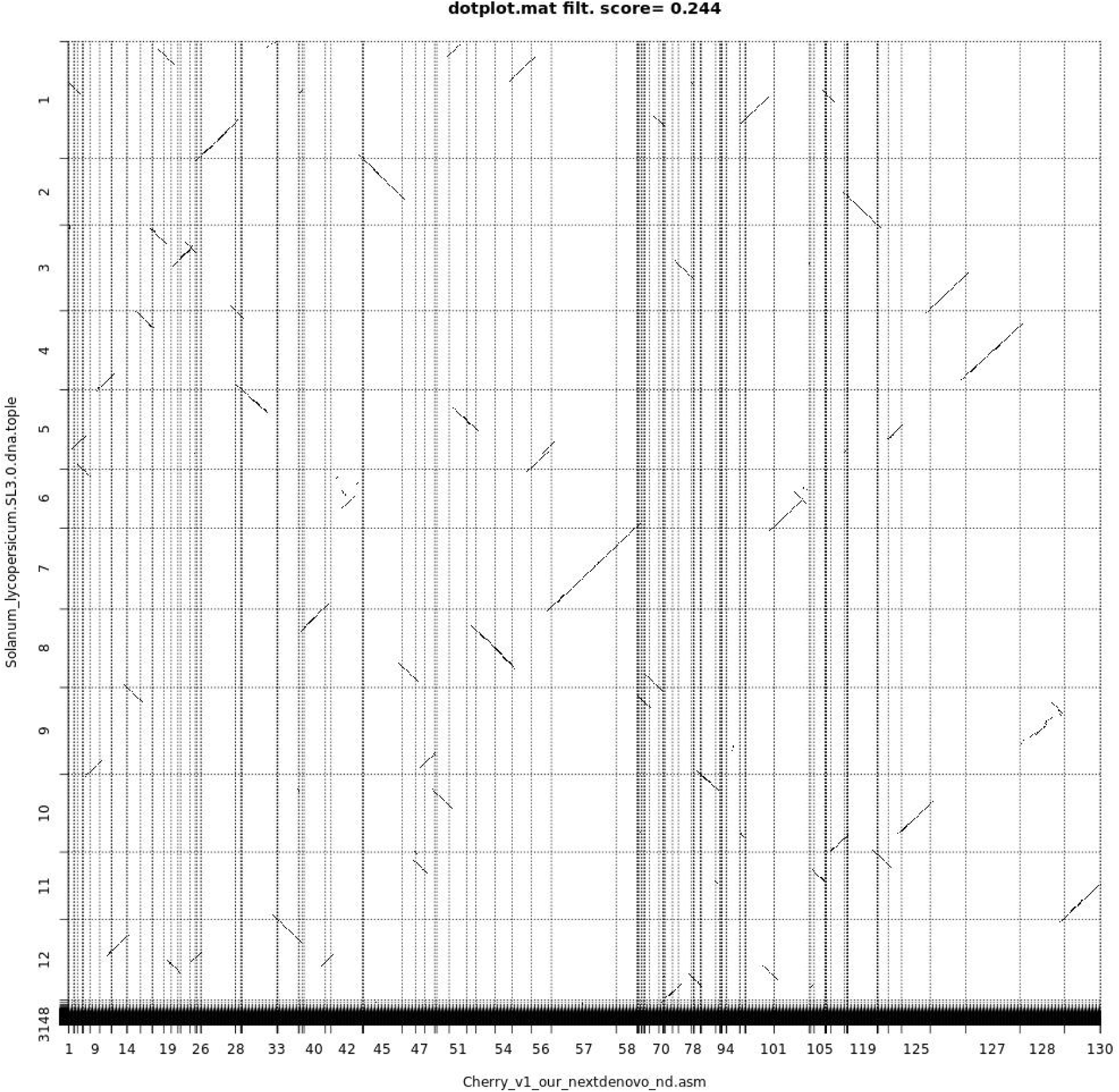
Dot plot for alignment of draft assembly contigs (X axis) and SL3.0 chromosomes (Y axis).

**Supplementary Figure S6.**
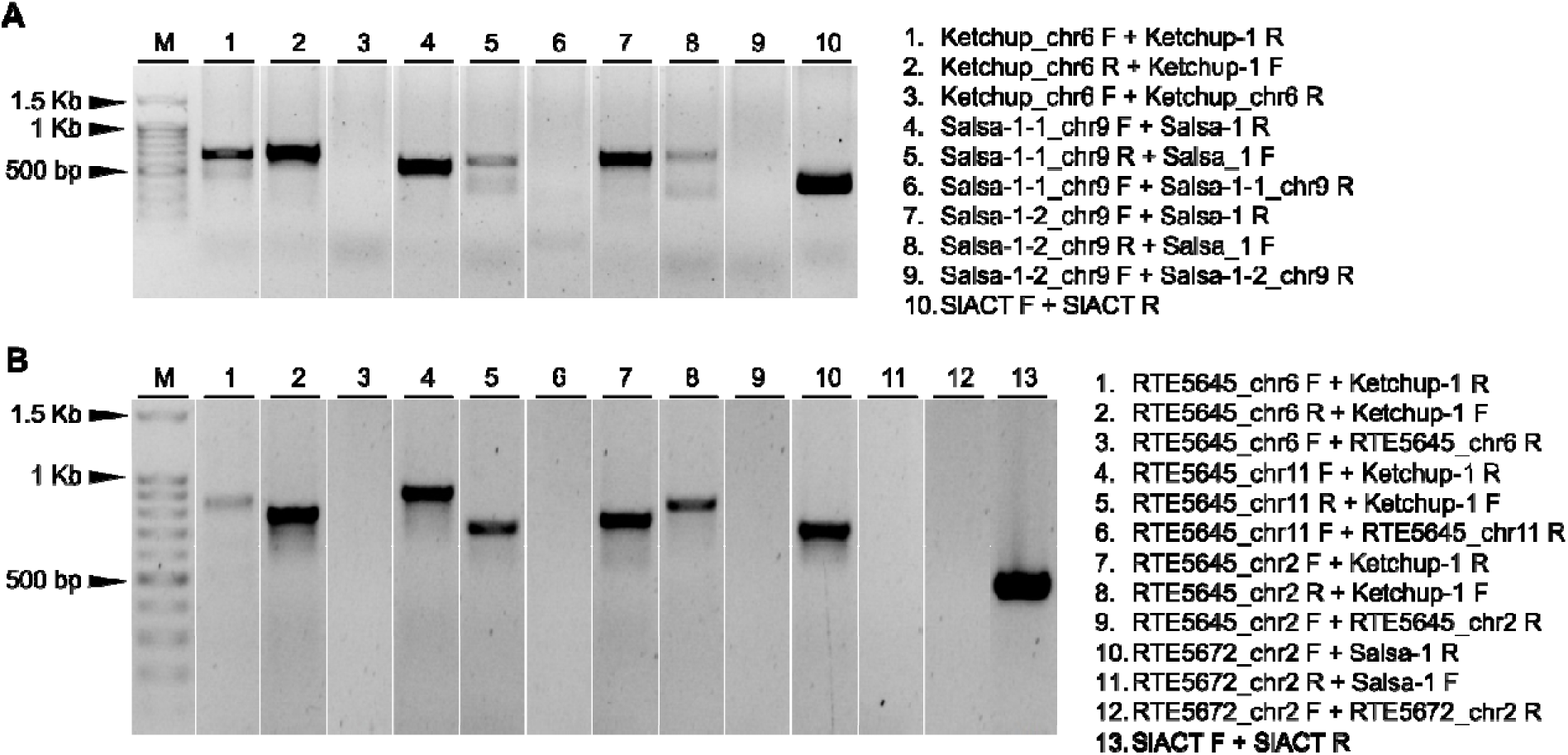
PCR validation of insertions of SP (A) and SL (B) *Ketchup and Salsa* elements TEIs in our tomato line.

