## Supplementary Figures for "Long-read sequencing of extrachromosomal circular DNA and genome assembly of a *Solanum lycopersicum* breeding line revealed active LTR retrotransposons originating from *S. peruvianum* L. introgressions"

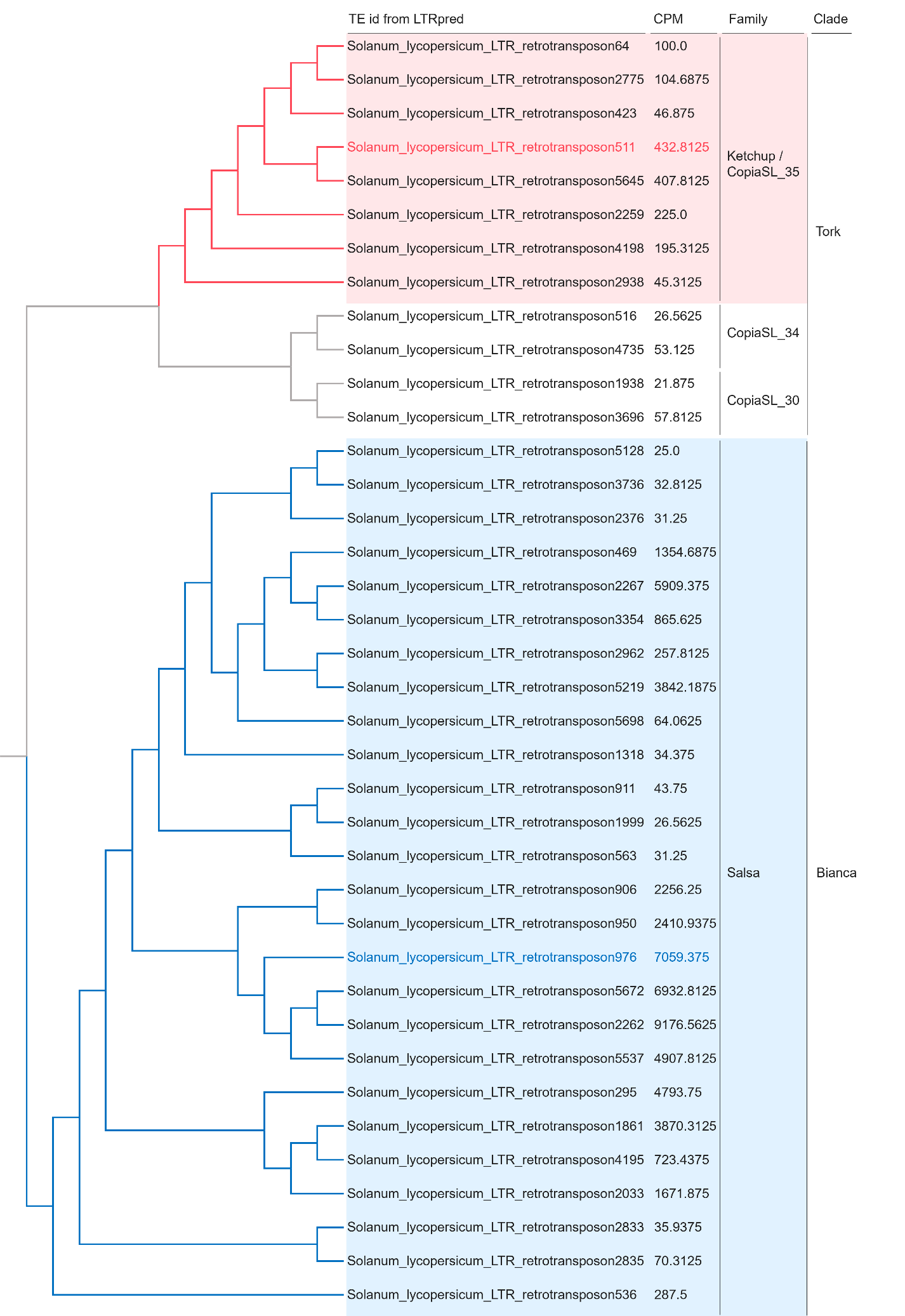

**Supplementary Figure S1.** Phylogenetic tree of full-length sequences of 38 RTEs. The red and blue colors correspond to the elements of the *Ketchup* and *Salsa* families, respectively.

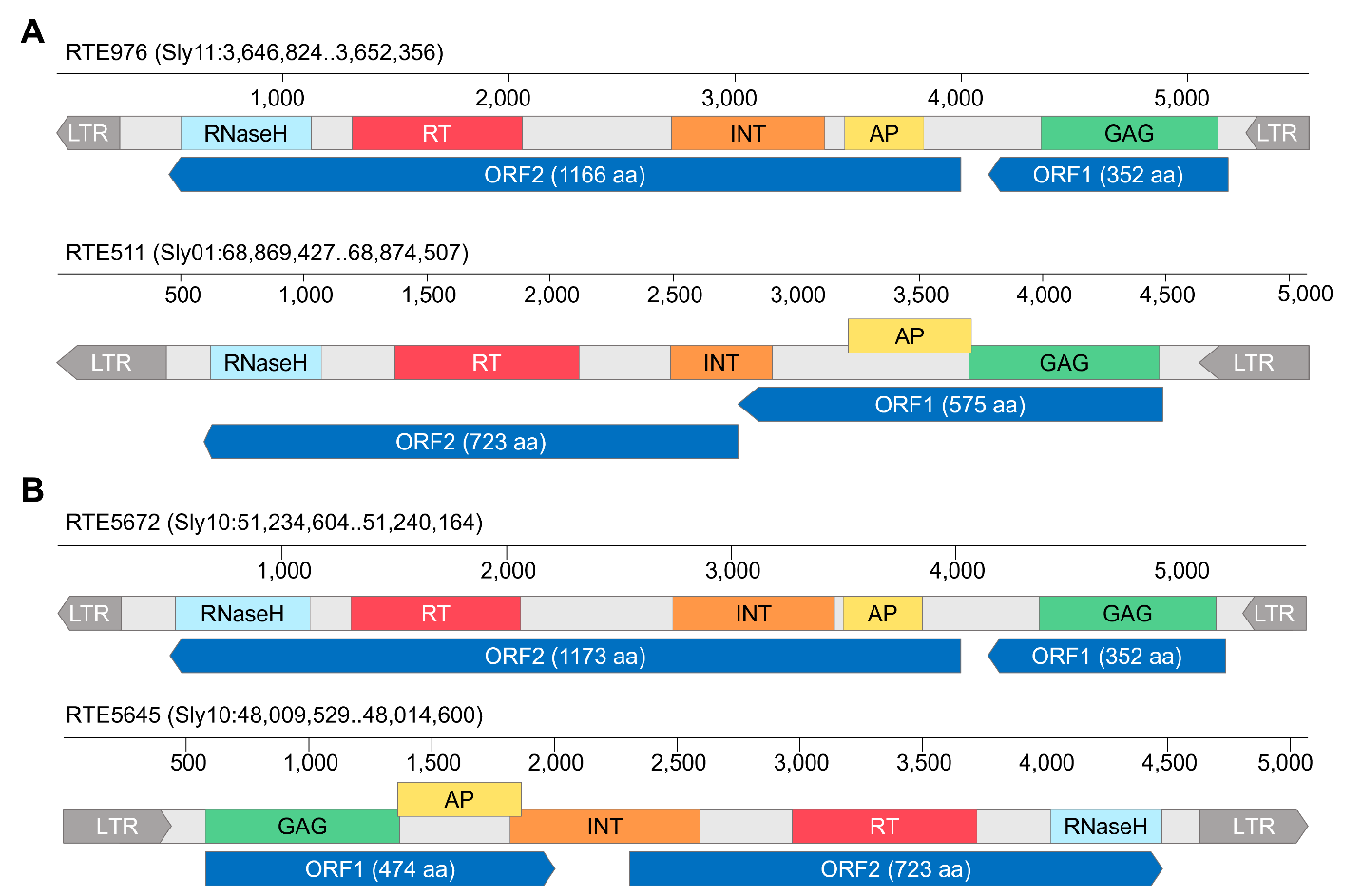

**Supplementary Figure S2.** Encoded domains and open reading frames (ORFs) for RTEs (A) RTE976 and RTE511, two SL elements belonging to the *Salsa* and *Ketchup* family respectively; (B) RTE5672 and RTE5645, two SL elements belonging to the *Salsa* and *Ketchup* family respectively.

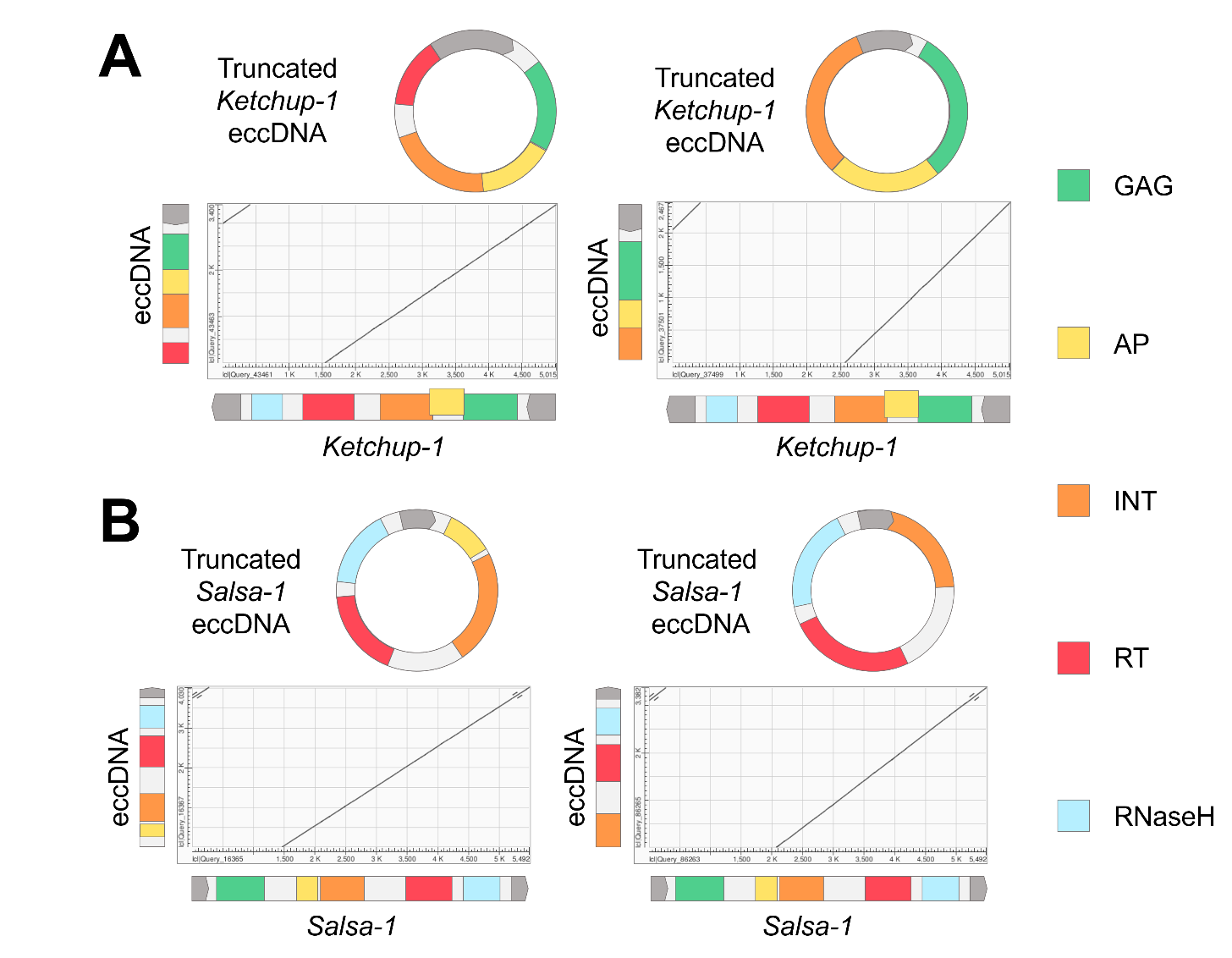

**Supplementary Figure S3.** Dot plots for alignment of truncated eccDNA sequences (Y axis) and full-length retrotransposon sequences (X axis) with domain annotation: (A) – *Ketchup-1*; (B) – *Salsa-1*.

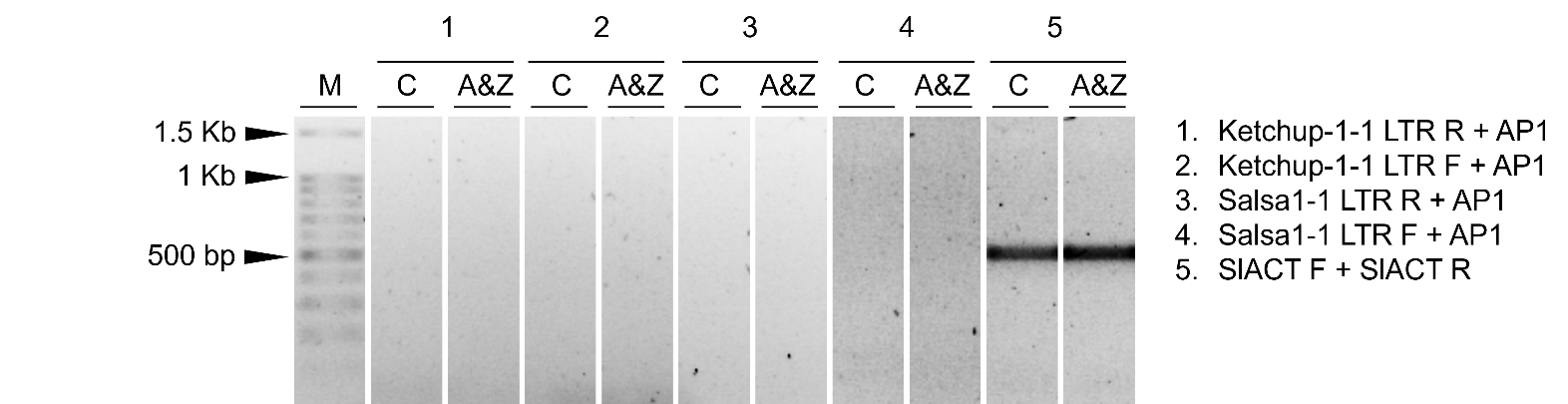

**Supplementary Figure S4.** PCR confirmation of the absence of eclDNA accumulation for *Ketchup-1-1* and *Salsa-1-1* elements in control (C) and relaxed TE control (A&Z) samples.

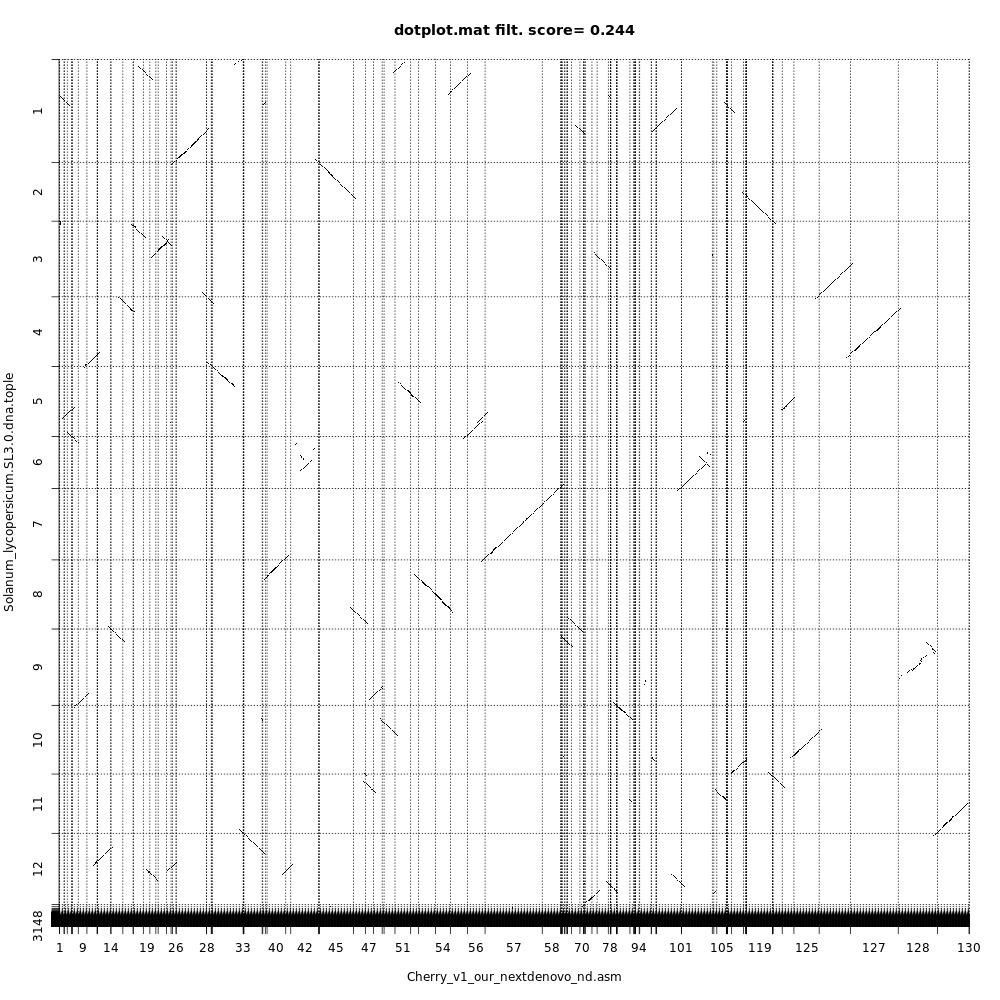

**Supplementary Figure S5.** Dot plot for alignment of draft assembly contigs (X axis) and SL3.0 chromosomes (Y axis).

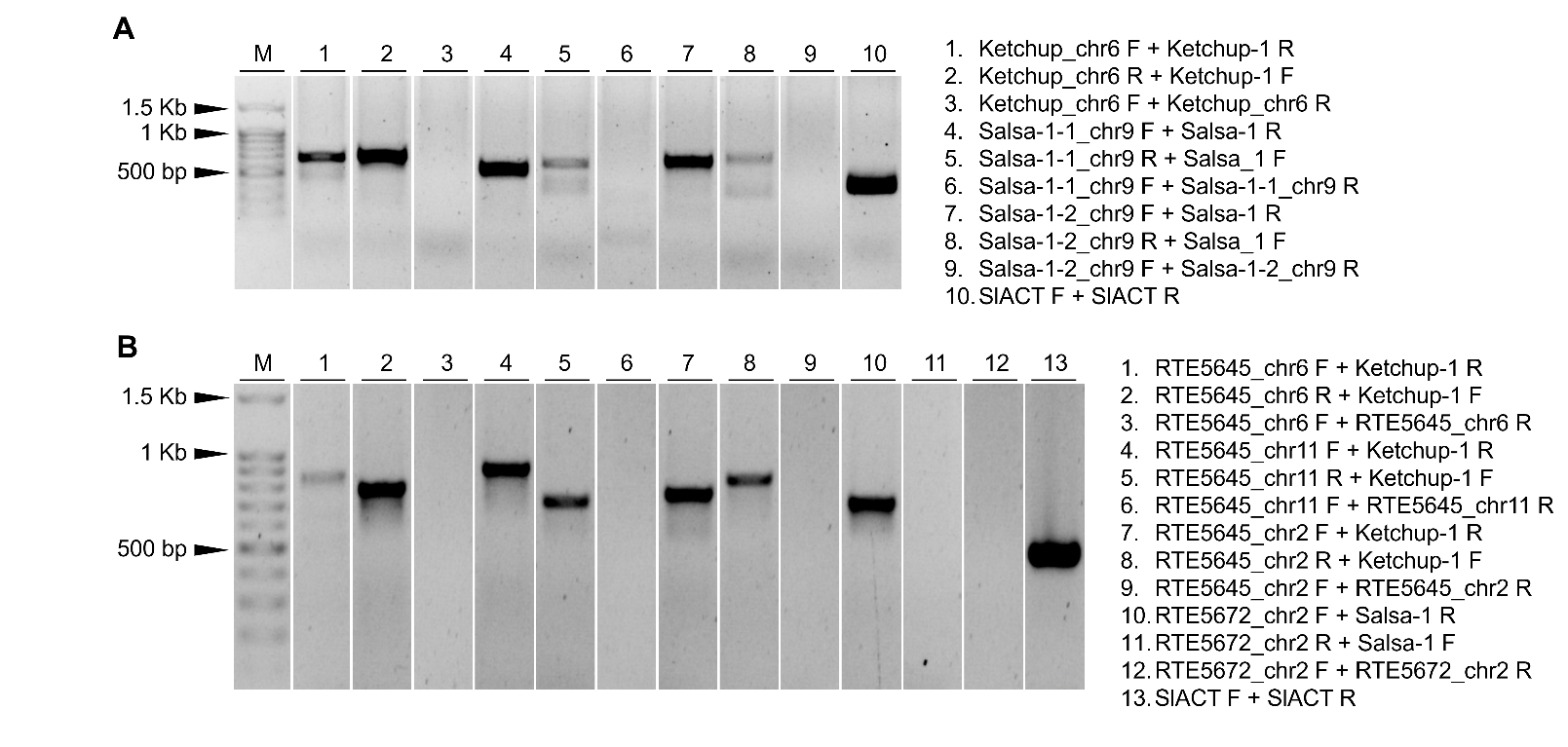

**Supplementary Figure S6.** PCR validation of insertions of SP (A) and SL (B) *Ketchup and Salsa* elements TEIs in our tomato line.
